# Systematic analysis of mouse genome reveals distinct evolutionary and functional properties among circadian and ultradian genes

**DOI:** 10.1101/197111

**Authors:** Stefano Castellana, Tommaso Mazza, Martin Kappler, Daniele Capocefalo, Nikolai Genov, Tommaso Biagini, Caterina Fusilli, Felix Scholkmann, Angela Relógio, John B. Hogenesch, Gianluigi Mazzoccoli

**Affiliations:** Bioinformatics Unit, IRCCS “Casa Sollievo della Sofferenza”, 71013 San Giovanni Rotondo (FG), Italy; Institute for Theoretical Biology (ITB), Charité–Universitätsmedizin and Humboldt University, Invalidenstraße 43, 10115 Berlin, Germany; Molekulares Krebsforschungszentrum (MKFZ), Charité-Universitätsmedizin Berlin, Augustenburger Platz 1, 13353 Berlin, Germany; Research Office for Complex Physical and Biological Systems (ROCoS), 8091Zurich, Switzerland; Department of Neonatology, University Hospital Zurich, University of Zurich, 8091 Zurich, Switzerland; Divisions of Human Genetics and Immunobiology, Cincinnati Children’s Hospital Medical Center, Cincinnati, OH, 45215, United States; Division of Internal Medicine and Chronobiology Unit, IRCCS “Casa Sollievo della Sofferenza”, 71013 San Giovanni Rotondo (FG), Italy

**Keywords:** clock, gene, evolution, rhythmicity, circadian, ultradian

## Abstract

In living organisms, biological clocks regulate 24 h (circadian) molecular, physiological, and behavioral rhythms to maintain homeostasis and synchrony with predictable environmental changes. Harmonics of these circadian rhythms having periods of 8 hours and 12 hours (ultradian) have been documented in several species. In mouse liver, harmonics of the 24-hour period of gene transcription hallmarked genes oscillating with a frequency two or three times faster than the circadian circuitry. Many of these harmonic transcripts enriched pathways regulating responses to environmental stress and coinciding preferentially with subjective dawn and dusk. We hypothesized that these stress anticipatory genes would be more evolutionarily conserved than background circadian and non-circadian genes. To investigate this issue, we performed broad computational analyses of genes/proteins oscillating at different frequency ranges across several species and showed that ultradian genes/proteins, especially those oscillating with a 12-hour periodicity, are more likely to be of ancient origin and essential in mice. In summary, our results show that genes with ultradian transcriptional patterns are more likely to be phylogenetically conserved and associated with the primeval and inevitable dawn/dusk transitions.

## INTRODUCTION

Recurring changes in environmental variables (i.e. temperature, atmospheric pressure, magnetism, ultraviolet radiation, humidity, food/water availability, etc.) greatly influence living organisms, impinging on processes and activities essential for individual and species survival. These include rest/wake and feeding/fasting cycles, hunting, courtship and mating behavior, *inter alia*.

Different molecular oscillatory mechanisms, consisting of endogenous non-transcriptional circuits as well as transcriptional/translational feedback loops, appeared within the Tree of Life as a result of convergent evolution, to manage energetic cycles driven by the solar light (Pittendrigh 1999 ^PMID:8466172^; Dunlap 1999 ^PMID:^ 9988221; Takahashi 2017 ^PMID: 27990019^). These biological clocks are common across species and phyla, allowing appropriate physiological/behavioral adaptation and rhythmicity to anticipate predictable environmental changes and providing survival advantage (Pittendrigh 1999 ^PMID8466172^; Dunlap 1999 ^PMID:^ 9988221; Takahashi 2017 ^PMID: 27990019^).

The most common and investigated biological rhythms have a roughly 24-hour periodicity and resonate with the daily switch from the bright light of day to the darkness of night. The mammalian biological clock drives the oscillatory expression of ~ 43% of all protein coding genes in an organ-specific manner (Bozek 2009 ^PMID: 19287494^; zhang 2014 ^PMID: 25349387^; Lehmann 2015 ^PMID:25945798^). In addition to the 24-hour periodicity, two clusters of genes cycling at the second (12-hour) and third (8-hour) harmonics of the circadian rhythm were identified through high temporal resolution analyses of the mouse transcriptome. These harmonics were found in intact mice and were lost *ex vivo* or under restricted feeding conditions (Hughes 2009 ^PMID:^ i93432o1 The repertoire0f rhythms observed in gene transcription could depend on binding strengths of the elements involved (genes and/or proteins) as well as transcription/translation dynamics involving degradation speed and temporal delays (Fuhr 2015 ^PMID:26288701^). As suggested, two-peak circadian oscillations in output gene expression may result from the interaction between transcription factors with definite circadian phase relationships and noncompetitive binding to the promoters of regulated genes (Westermark 2013 ^PMID: 23583178^) or through an oscillatory incoherent feed-forward loop (Martins 2016: 28007935) Despite the pervasiveness of biological clocks among species, evolutionary and functional properties of oscillating genes remain largely unexplored.

Among higher organisms, the mouse is the favorite animal model for studying biological rhythms from a genomic and molecular point of view. Oscillating genes have been observed to cluster together along the mouse genome, to be longer and to have more alternative transcripts with respect to non-oscillating genes (Zhang 2014 ^PMID:25349387^). Regulation of oscillating genes expression has been also extensively studied in mouse: three main cis-regulatory elements are located in the proximity of gene promoters (E-box, D-box, Ror/Rev-Erb response elements) and allow their time-regulated expression [Takahashi 2017 ^PMID: 27990019^].

Furthermore, insights into the physiological role of ultradian oscillating genes have been published, suggesting the importance of the ultradian expression of genes involved in metabolism regulation, feeding behavior and response to dawn/dusk transition-related stress situations (Hughes et al.2009 ^PMID:19343201^).. Another intriguing aspect of oscillating gene regulation regards codon usage and translation: recent experimental evidences found that key circadian genes within simple organisms (i.e., *Neurospora crassa, Synechococcus elongatus, Drosophila melanogaster)* exhibit an unexpected non-optimized codon usage (Zhou et al., 2013; Zhou et al., 2015; Xu et al., 2013; Fu et al., 2016).

Thus, in this study we asked ourselves how and when genes with the three different oscillatory patterns originated along the Tree of Life. Furthermore, inspired by the abovementioned data regarding codon usage impact on circadian gene expression, we explored Synonymous Codon Usage for the whole set of mouse oscillating genes, by partitioning the gene set in the three period classes and comparing them to the global genetic background. Moreover, we inspected upstream genomic sequences within the mouse genome trying to evidence Transcription Factor Binding Sites (TFBS), other than known *cis*-regulatory elements that could provide additional insights into the regulation of oscillating genes.

We also carried out a comparative analysis of functional annotations for the 24h, 12h and 8h oscillating genes and evaluated the impact of these genes on organism viability (gene essentiality). Function and evolutionary conservation of ultradian mouse genes would suggest that such genes have played a crucial role during life evolution, maybe determining a proper response to environmental stress conditions. We hypothesized that at various levels of biological organization their regulation was adapted in order to finely manage stress conditions dictated by recurrent transitions along the day, like the dawn/dusk transitions.

## RESULTS

We investigated different aspects of mouse genomics in order to better understand the relationships between evolutionary origin, regulation and function of the different oscillating gene subsets. Our reference mouse gene lists were taken from a single publicly available data set (Hughes 2009 ^PMID: 19343201^). Different valuable although not fully comparable methodological approaches have been recently used in order to detect circadian and ultradian signals within transcriptomes (e.g., Zhu 2017 ^28591634^; Van der Veen 2017 ^27871062^). The results of our well-established bioinformatics approach partly overlap with ultradian gene sets identified by these studies, which define different criteria to identify harmonics of circadian rhythmicity in gene expression.

### Phylogenetic origin of cycling genes

We operated a systematic inspection of oscillating gene trees retrieved from the PhylomeDB, NCBI HomoloGene, Phyletic age resources [Huerta-Cepas 2014 ^PMID: 24275491^; NCBI Reference Coordinators 2017 PMiD: 27899561. 2017 ^PMID:27799467^].We considered the oldest phylogenetic node for each gene and those genes that originated early during evolution (Opisthokonts) were labeled as “ancient”. Genes that presumably arose after Chordate radiation were labeled as “recent” (cf., Table 1). Raw data obtained from PhylomeDB, NCBI HomoloGene and Phyletic age are reported in Supplementary Table 1.

**Table 1.** Evolutionary origin of 8h, 12h and 24h oscillating genes sorted out of the global mouse transcriptome, according to PhylomeDB, NCBI HomoloGene and Phyletic age. For each taxon, the number of genes with putative origin in that taxon are shown.

| **PhylomeDB** | **8h** | **12h** | **24h** | **All genes** |
| --- | --- | --- | --- | --- |
| **Taxonomic groups** | (n=43/56)<br>% (n) | (n=164/202)<br>% (n) | (n=1580/2054)<br>% (n) | (n=15443)<br>% (n) |
| *Prokaryota* | 9.3 (4) | 9.8 (16) | 4.8 (76) | 5.7 (879) |
| *Eukaryota* | 27.9 (12) | 52.4 (86) | 30.4 (481) | 29.2 (4516) |
| *Opisthokonts* | 37.2 (16) | 20.1 (33) | 20 (316) | 21.7 (3357) |
| *"Ancient" genes* | 74.4 (32) | 82.3 (135) | 55.2 (873) | 56.6 (8752) |
| *Chordate* | - | - | - | - |
| *Teleostomi* | 20.9 (9) | 12.2 (20) | 20.2 (319) | 22.8 (3525) |
| *Tetrapoda* | - | - | - | - |
| *Amniota* | 0 (0) | 2.4 (4) | 11.3 (179) | 8.8 (1346) |
| *Mammalia* | - | - | - | - |
| *Eutheria* | 4.7 (2) | 3 (5) | 13.3 (209) | 11.8 (1820) |
| *"Recent" genes* | 25.6 (11) | 17.7 (29) | 44.8 (707) | 43.4 (6691) |
| **NCBI HomoloGene** | **8h** | **12h** | **24h** | **All genes** |
| **Taxonomic groups** | (n=49/56)<br>% (n) | (n=183/202)<br>% (n) | (n=1474/2054)<br>% (n) | (n=19032)<br>% (n) |
| *Eukaryota* | 20.4 (10) | 35.5 (65) | 15.9 (234) | 13.1 (2500) |
| *Opisthokonts* | 10.2 (5) | 2.7 (5) | 3.7 (54) | 3.1 (587) |
| *"Ancient" genes* | 30.6 (15) | 38.2 (70) | 19.6 (288) | 16.2 (3087) |
| *Metazoa* | - | - | - | - |
| *Bilateria* | 22.4 (11) | 25.7 (47) | 17.6 (259) | 15.1 (2883) |
| *Vertebrata* | - | - | - | - |
| *Euteleostomi* | 30.6 (15) | 25.1 (46) | 40.2 (592) | 34.6 (6579) |
| *Tetrapoda* | 4.2 (2) | 7.2 (13) | 8.6 (127) | 7.7 (1459) |
| *Amniota* | - | 1.6 (3) | 3 (45) | 3.5 (657) |
| *Mammalia* | - | - | - | - |
| *Boreoeutheria* | 12.2 (6) | 2.2 (4) | 10 (148) | 14.8 (2825) |
| *Euarchontoglires* | - | - | 0.9 (14) | 1.9 (357) |
| *Murinae* | - | - | 0.1 (1) | 5.2 (996) |
| *exclusive to Mus musculus* | - | - |  | 1 (189) |
| *"Recent" genes* | 69.4 (34) | 61.8 (113) | 80.4 (1186) | 83.7 (15945) |
| **Phyletic age** | **8h** | **12h** | **24h** | **All genes** |
| **Taxonomic groups** | (n=52/56)<br>% (n) | (n=189/202)<br>% (n) | (n=1882/2054)<br>% (n) | (n=22544)<br>% (n) |
| *Eukaryota* | 36.5 (19) | 41.3 (78) | 25.1 (472) | 23.2 (5262) |
| *Opisthokonts* | 1.9 (1) | 8.5 (16) | 6.6 (124) | 54.2 (1227) |
| *"Ancient" genes* | 38.5 (20) | 49.7 (94) | 31.7 (596) | 28.8 (6489) |
| *Metazoa* | 32.7 (17) | 28 (53) | 25.9 (488) | 2.2 (5351) |
| *Chordate* | 21.1 (11) | 13.2 (25) | 24.2 (455) | 23.7 (4753) |
| *Mammalia* | 3.8 (2) | 3.7 (7) | 9 (169) | 21.1 (2011) |
| *exclusive to Mus musculus* | 3.8 (2) | 5.3 (10) | 9.2 (174) | 17.5 (3940) |
| *"Recent" genes* | 61.5 (32) | 50.3 (95) | 68.3 (1286) | 71.2 (16055) |
| All genes refers to the global mouse genetic background |  |  |  |  |

From PhylomeDB, we were able to retrieve phylogenetic information for 15443 genes out of the global mouse transcriptome and for 1787 out of 2312 oscillating mouse genes. Genes oscillating with a 8-hour and 12-hour periodicity were significantly enriched in “ancient” genes with respect to the global mouse transcriptome (Fisher’s Exact Test for Count Data; respectively: *p* = 0.01 and *p* = 3.58e^−12^; 95% C.I. 1.21-Inf, 2.51-Inf; Odds Ratios 2.22, 3.55). No enrichment was found for mouse genes oscillating with a 24-hour period. Besides, 8h and 12h oscillating genes were enriched in “ancient” genes with respect to the 24h oscillating ones (respectively: *p* = 0.008 and *p* = 2.37e^−12^; 95% C.I.: 1.26-Inf and 2.63-Inf; Odds Ratios: 2.35 and 3.76).

We then analyzed homologous genes in NCBI Homologene, with particular focus on *Mus musculus* species group. We retrieved homologous accession numbers, gene symbols and conservation data from the summary output, mapped oscillatory genes, and calculated the relative proportions. We retrieved 1706 oscillating genes out of the global mouse transcriptome (19032 genes), split them according to the “ancient” and “recent” classes, and calculated asymmetries among counts. The 12h oscillating gene subset was strongly enriched in “ancient” genes (*p* = 9.72e^−13^; 95% C.I. 2.45-Inf; Odds Ratio 3.19). In this analysis, circadian genes and 8h oscillating genes were slightly enriched in “ancient” genes with respect to the global mouse transcriptome *(p =* 0.01; 95% C.I. 1.11-Inf; O.R 1.25; *p* = 0.0091; 95% C.I. 1.28-Inf; Odds Ratio 2.27). Statistically significant differences emerged comparing 12h and 24h oscillating genes (*p* = 3.57e^−8^; 95% C.l: 1.91-Inf; O.R. 2.55) and, less significantly, comparing 8h and 24h oscillating genes (*p* = 0.046, 95% C.I. 1.01-Inf, O.R. 1.81).

Finally, we analyzed oscillatory genes with the Phyletic age dataset. Genes oscillating with a 12-hour periodicity were still enriched in “ancient” genes with respect to the global mouse transcriptome (Fisher’s Exact Test *p =* 1.28e^−9^; 95% C.I. 1.9-Inf; Odds Ratio 2.44) and compared to the 24h oscillating gene subset *(p =* 7.15e ^−7^; 95% C.I. 1.63-Inf; Odds Ratio 2.13). The 8h oscillating gene subset did not differ from the other subsets and the global mouse transcriptome. Circadian genes were also significantly enriched in “ancient” genes when compared to the global mouse transcriptome (*p* = 0.004; 95%C.I.1.05-Inf; Odds Ratio 1.14).

### Protein age estimation and enrichment analysis

Proteins found in different living species arose at specific evolutionary times and their evolutionary origins provide information about function and interactions. To further explore the evolutionary conservation of the three oscillating gene subsets at the protein level, we determined their age using the ProteinHistorian tool [Capra 2012 ^PMID: 22761559^]. Each comparison was performed with respect to the average age of the genomic background (553 million years). Our data showed that the 8h and the 12h oscillating gene subsets have a similar average computed age of 1117 million years and of 1143 million years, respectively, while the 24h oscillating gene subset was found to be less ancient with an average age of 973 million years (Figure 1 A-C). Interestingly, the three oscillating gene subsets were more ancient than the genomic background, and can be traced back to very ancient branches of the Tree of Life. In the case of the 8h oscillating gene subset, the bins containing the highest number of genes are located at the Euteleostomi and Eukaryota branches (Figure 1A). As for the 12h oscillating gene subset, the bins containing the highest number of genes are located at the Eukaryota branch (Figure 1B). As for the 24h oscillating gene subset, we found three large bins at Euteleostomi, Eukaryota and Deuterostomia branches (Figure 1C). Nevertheless, given the larger size of the 24h oscillating gene subset, as compared to the genes oscillating with 12-hour and 8-hour periodicity, it is conceivable that smaller sets of ancient genes exist, but that these are masked by the large size of the background set of genes. The fact that all the three oscillating gene subsets contain a large bin at the Eukaryota level, hints that most of the oscillating genes developed early during the evolution of life on Earth and have been conserved over a long period of evolutionary time, suggesting that they may play an essential function in many organisms.

**Figure 1.**
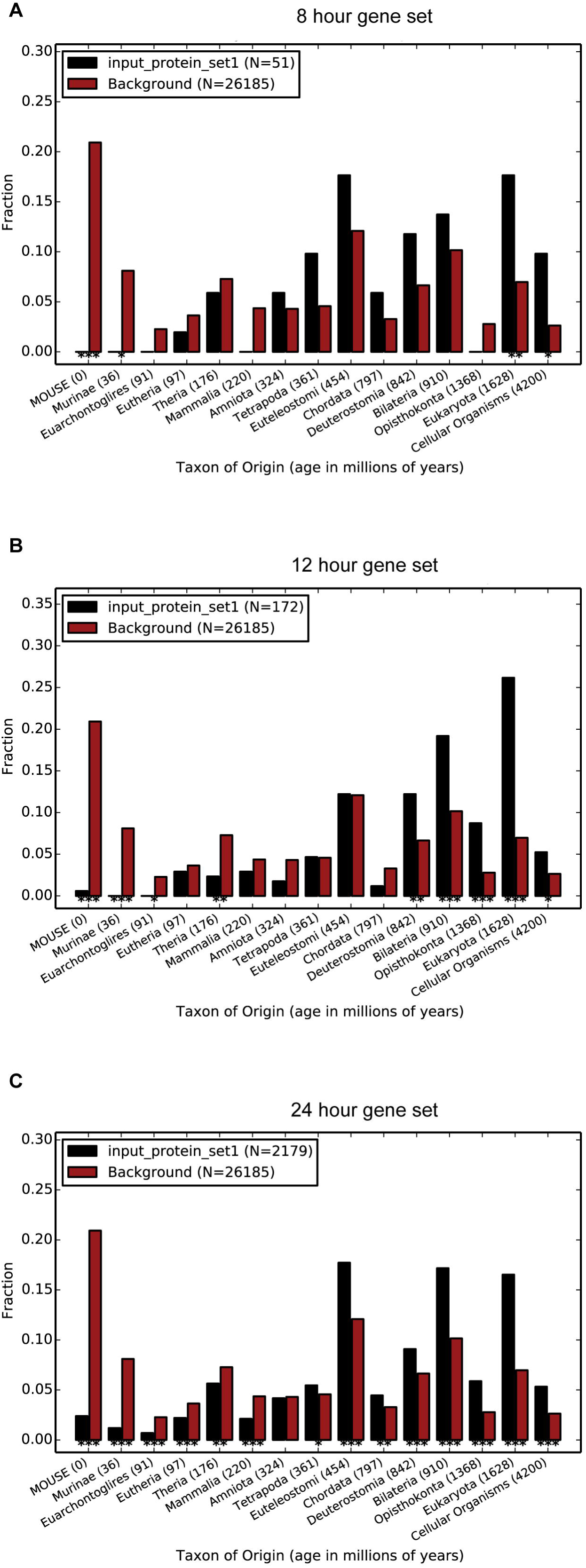
The evolutionary age of the oscillating gene subsets differs significantly. The 12h oscillating gene subset shows the highest average age. The 8h (A) oscillating gene subset has an average age of 1117 million years, the 12h (B) oscillating gene subset has an average age of 1143 million years and the 24h (C) oscillating gene subset has an average age of 973 million years. All the three subsets are significantly older than the genomic background, having an average age of 553 million years. Since the gene subsets are derived from experiments performed with modern mice, the age of the mouse has been set to 0 million years on the x-axis. The remaining ages refer to the emergence of the corresponding branches of the Tree of Life. Data for the computed evolutionary age of the oscillating gene subsets were gathered by the Protein Historian tool (http://lighthouse.ucsf.edu/ProteinHistorian/).

### Essentiality of oscillating genes

Genes that are necessary for an organism’s survival are defined as “essential”. Gene essentiality refers to the fact that such genes play a fundamental role so that their dysfunction cannot be compensated by other genes or molecular mechanisms. Essential genes are then, theoretically, more likely to be of ancient origin because they constitute the “minimal” necessary genetic repertoire in order for a living being to survive in its environment.

By consulting the OGEEv2 databank (http://ogee.medgenius.info/downloads/) (Chen 2017 ^PMID: 27799467^); we retrieved information for mouse genome essentiality, by focusing on oscillating gene subsets (Table 2).

**Table 2.** Number and proportion of essential/non-essential oscillating mouse genes according to the OGEEv2 database

|  |  |  |  |  |
| --- | --- | --- | --- | --- |
| *Essentiality* | *8h* | *12h* | *24h* | *All genes* |
| Essential | 15 (43%) | 63 (68.5%) | 392 (47%) | 4341 (48%) |
| Non-essential | 20 (57%) | 29 (31.5%) | 446 (53%) | 4700 (52%) |
| Total genes retrieved | 35 (100%) | 92 (100%) | 838 (100%) | 9041 (100%) |
| All genes refers to the global mouse genetic background |  |  |  |  |

We extracted and focused our analysis on 9041 mouse genes with some experimental indication of essentiality/non-essentiality. The 68.5% of genes oscillating with 12-hour periodicity were categorized as essential with respect to the set of all genes (Fisher’s Exact Test for Count Data; *p* = 6.305^e-5^, C.I.:1.59-Inf, O.R.:2.35). No significant disproportion was found for the 8h and 24h oscillating gene subsets. These data corroborate the hypothesis that the 12h oscillating genes are critical components of the eukaryotic cell, in order that they tend to be evolutionary conserved and essential.

### Synonymous codon usage in oscillating genes

The different patterns of synonymous codon usage (SCU) within and among genomes have been generally related to efficient translation or accurate protein folding. Although intriguing findings emerged for important circadian genes within basal species *(Neurospora crassa, Synecocchus elongates, Drosophila melanogaster)*, no systematic inspection of codon usage has been carried out in complex species, such as *Mus Musculus*.

First, we calculated the Effective Number of Codons (ENC) and the Codon Adaptation Index (CAI) for the total pool of mouse protein-coding genes (26780 common sequences between the CodonW and Jcat outputs) and then evaluated potential differences among oscillatory gene subsets and the genetic background.

Given that preferentiality of SCU is a tradeoff between gene-specific mutational bias and purifying selection, we performed a further analysis by removing genes with extreme compositional properties. We considered the GC content of the third codon positions (‘GC3s’) as an estimate of the mutational pressure within genes: thus, we focused only on genes within the lowest and highest quartile (0.48 < GC3s < 0.66). Table 3 resumes comparative tests for the two indexes and the 3 oscillatory gene classes (‘n’ indicates the number of the resulting sequences after GC correction). Genes that oscillate with 12-hour periodicity generally have non-optimized SCU (higher ENC and smaller CAI with respect to the 13323 background sequences), while the SCU of the other gene subsets was found to be similar to the general scenario. Raw outputs of CodonW and JCat tools are presented in Supplementary Table 3 and original output of CodonW and JCat in, respectively, Supplementary File 1 and 2.

**Table 3.**
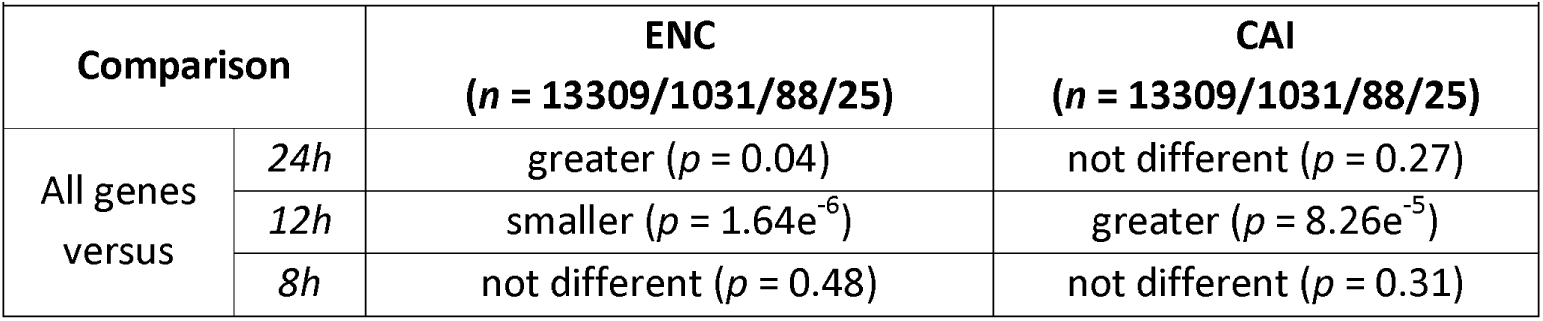
Results of Wilcoxon Tests for ENC/CAI distributions of 24h, 12h and 8h oscillating gene subsets against the whole mouse genome; “n” represent s the size of each distribution

Figure 2 shows box plots rendering a descriptive view of the ENC and CAI value distribution among the oscillating gene subsets with different oscillation periodicity

**Figure 2.**
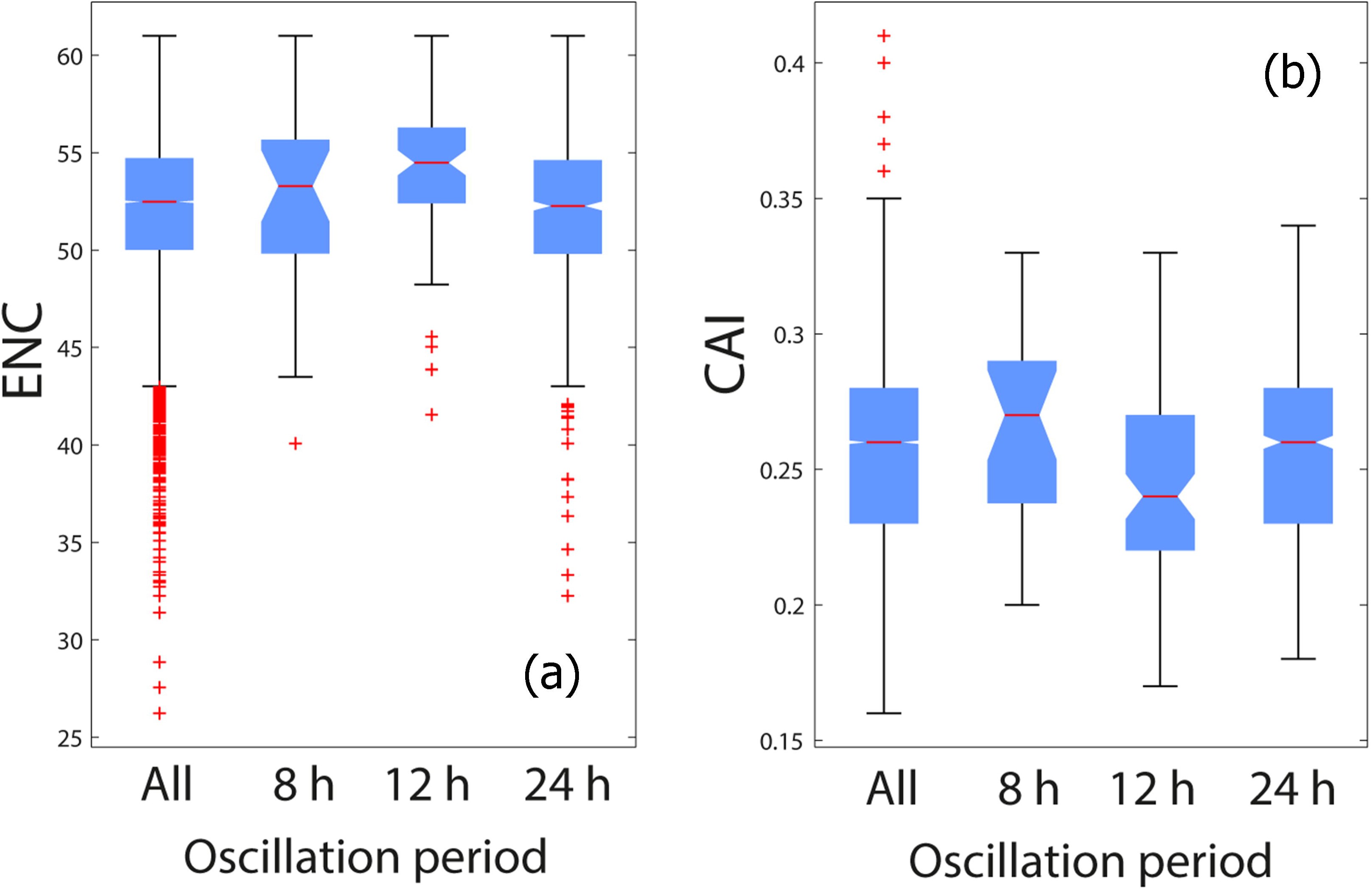
A) Box plots of ENC values calculated for all selected mouse transcript sequences and the three oscillating gene subsets. Selection was carried out according to the GC content (see Methods for details). B) Box plots of CAI values calculated for all selected mouse transcript sequences and the three oscillating gene subsets. Selection was carried out according to the GC content (see Methods for details).

### Functional differences among *oscillating gene subsets*

We surveyed the functional impact of mouse genes oscillating with 8-hour, 12-hour and 24-hour periods by performing functional (pathway, gene ontology) enrichment analyses through the use of Ingenuity Pathway Analysis (IPA) (QIAGEN Inc., Ingenuity Pathway Analysis) (Supplementary Table 5).

**Supplementary Table 5.**
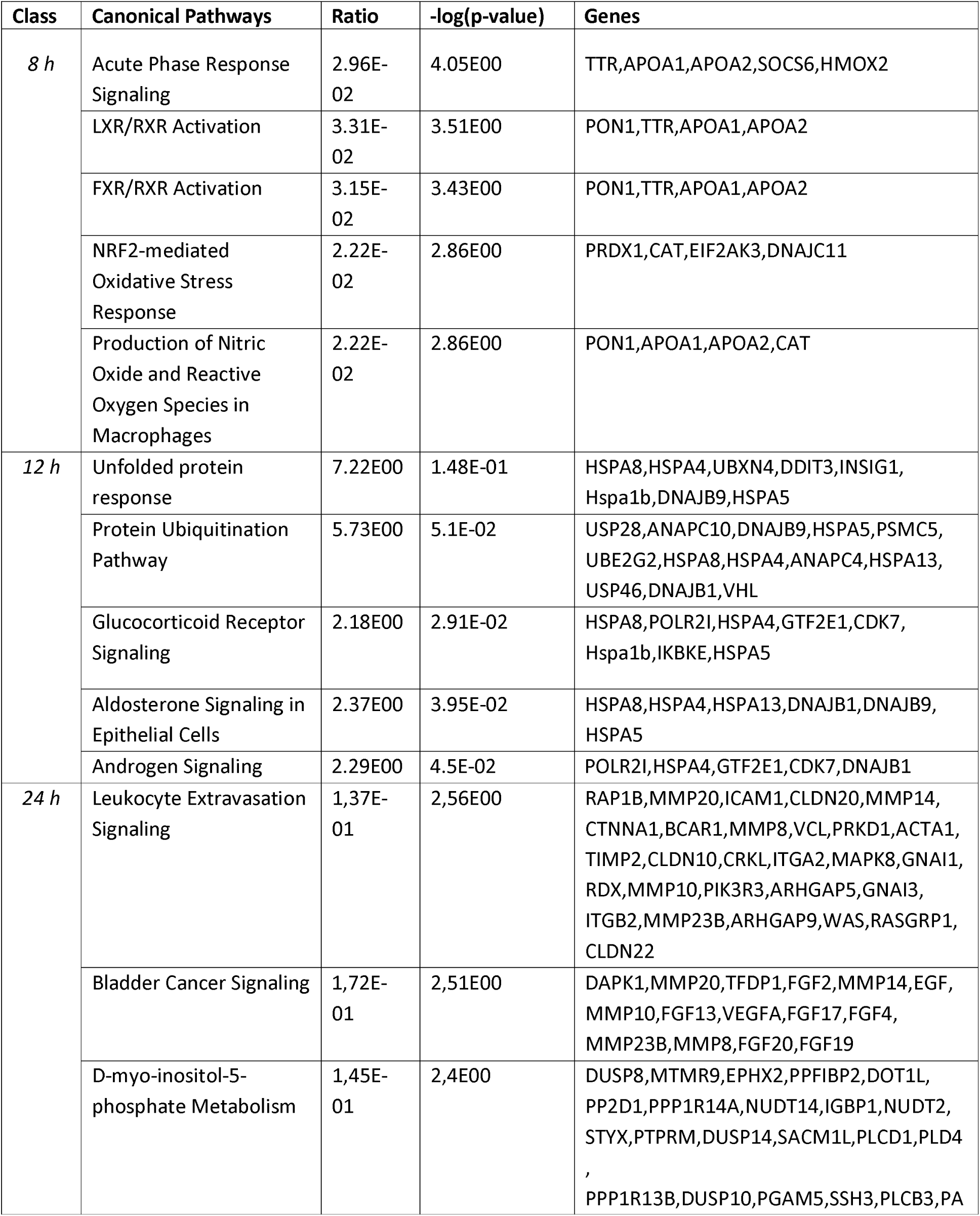
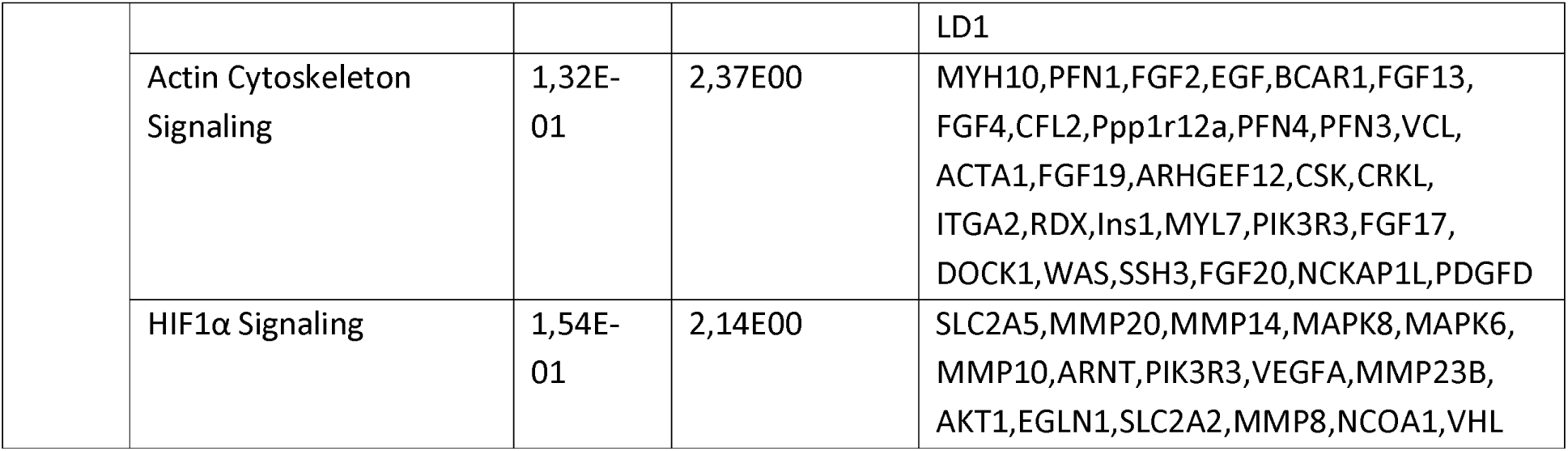
Top five enriched pathways for the three oscillating gene subsets

Furthermore, we performed a comparative analysis, evidencing differentially enriched pathways among the three oscillating gene subsets, although only the top-ranking ones are reported (Supplementary Table 6).

**Supplementary Table 6.** Comparative analysis based on -log(p-value) of enriched pathways among the three oscillating gene subsets

|  |  |  |  |
| --- | --- | --- | --- |
| *IPA Canonical Pathway* | *24h* | *12h* | *8h* |
| Protein Ubiquitination Pathway | 0.52 | **5.64** | 0.33 |
| Acute Phase Response Signaling | 0.43 | 0.37 | **5.49** |
| Unfolded protein response | 0.00 | **5.20** | 0.92 |
| NRF2-mediated Oxidative Stress Response | 1.58 | 1.17 | **3.05** |
| Production of Nitric Oxide and Reactive Oxygen Species in Macrophages | 0.84 | 0.68 | **2.99** |
| Complement System | **2.08** | 0.00 | **2.43** |
| FXR/RXR Activation | 0.72 | 0.00 | **3.49** |
| LXR/RXR Activation | 0.00 | 0.00 | **3.60** |
| Endoplasmic Reticulum Stress Pathway | 0.30 | **1.87** | 1.31 |
| Atherosclerosis Signaling | 0.94 | 0.00 | **2.48** |

The most overrepresented pathways for genes oscillating with 8-hour periodicity are related to sterol and fatty acids metabolism as well as oxidative stress, while a significant portion of 12h oscillating genes is involved in protein folding/ubiquitination and hormone signaling. Regarding 24h oscillating genes, many functions/pathways resulted to be significantly enriched and among these the category ‘circadian rhythm’ was ranking among the statistically significant pathways in our gene expression analysis in mouse liver (For more information see Supplementary Table 4).

### Enrichment analysis of TFBS within oscillating genes

We explored the genomic context of ultradian genes in search of over-represented binding sites. The possible presence of additional cis-regulatory elements that are specific for 8h and 12h oscillating gene subsets beyond well-established cis-acting elements associated with 24h oscillating gene expression could influence their different oscillatory pattern with respect to the circadian genes.

The implementation of AME tool on oscillating vs. non-oscillating genes upstream sequences revealed very interesting results, particularly with regard to the subset of 202 genes oscillating with a 12-hour period. We were able to retrieve the following enriched motifs (adjusted p-value<0.05) for the comparisons with five distinct random datasets (12, 13, 10, 14, 9; see methods). These pools greatly overlap within comparative analyses, with five common inferred TFBS (Zfp1_primary, Zfpl_secondary, XBP1.C, MBD2.B, GABPA.A) frequently resulting as over-represented. A significant enrichment for these five elements was also observed by considering upstream sequences of genes oscillating with a 12-hour periodicity and upstream sequences of 24h oscillating genes as control dataset.

No significantly enriched motif was detected when comparing upstream sequences of 8h oscillating genes with the random datasets and for 4 out of 5 comparisons of upstream sequences of genes oscillating with a period of 8 hours and upstream sequences of 24h oscillating genes as control dataset.

Two motifs (ERR3.B, E4F1.D) resulted to be enriched within upstream regions of genes oscillating with a 24-hour periodicity with respect to a single random sample of 1500 upstream sequences.

Supplementary Table 7 lists the top ranking motifs with significant enrichment in almost all comparative analyses; showed p-values are specific for the first comparison, although they result very low for all the performed comparative analyses (command lines and raw output in Supplementary File S3).

**Supplementary Table 7.**
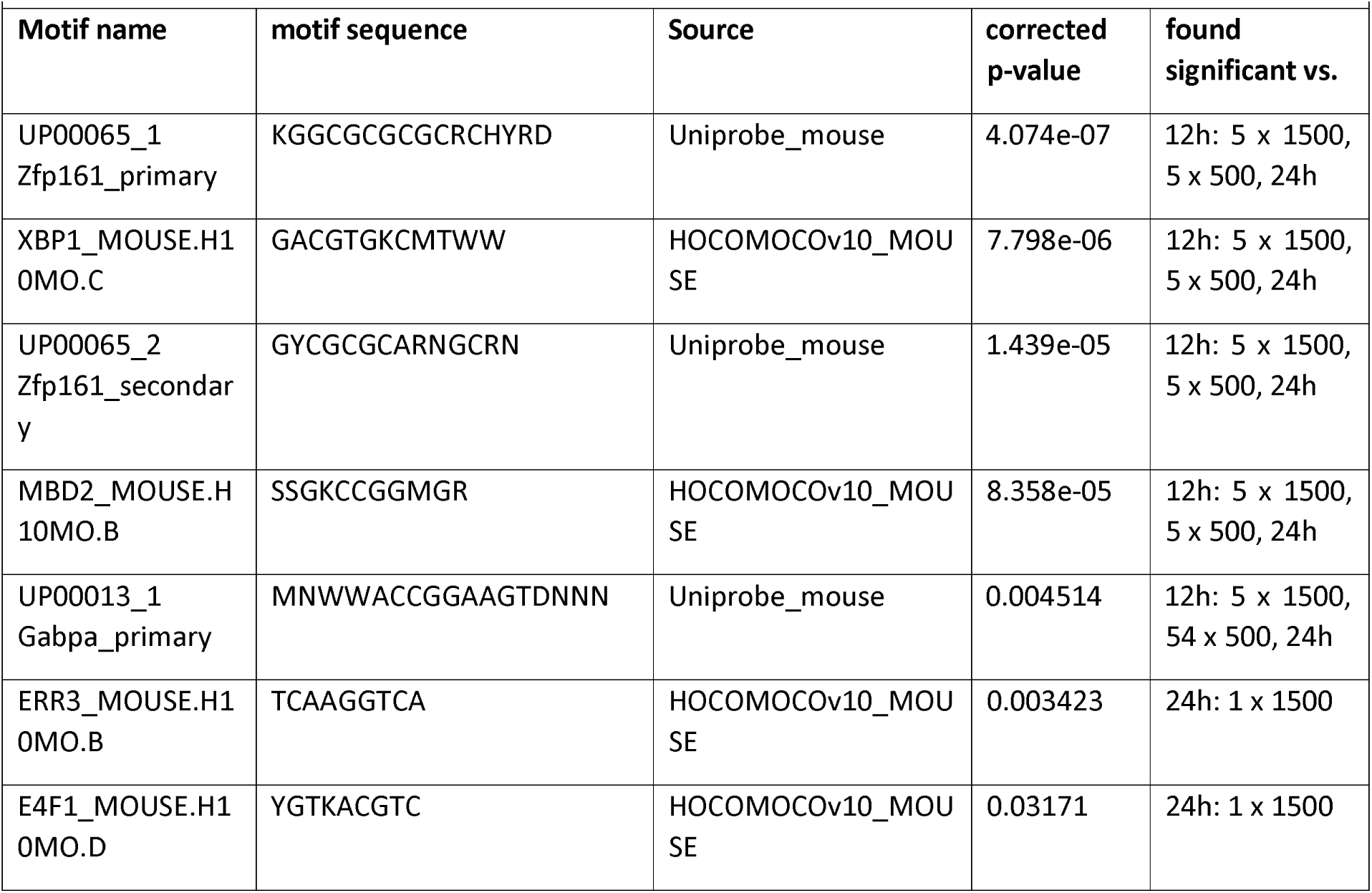
Summary of the top ranking over-represented Transcription Factor Binding Sites, as estimated by AME (MEME suite) comparative analyses

## DISCUSSION

The basic role played by biological timing systems is reflected by the fact that molecular oscillators tick in the three domains of life and oscillating genes have been discovered in phylogenetically distant species, ranging from cyanobacteria to humans. The cycling period of biological rhythms spans from minutes to years (Mazzoccoli 2016 ^PMID: 26906327^), but perhaps the largest class is attuned with a 24-hour rhythmicity dictated by Earth’s rotation around its axis. Different astronomical and geological factors affected Earth’s rotation during its history. While estimates of the rotation period are imprecise, when early life appeared, day length was significantly shorter than today. Interestingly, in relation to the highly metamorphosed nature of the oldest sedimentary rocks, there are no confirmed microfossils older than 3.5 billion years on Earth, but proofs support biological activity in submarine-hydrothermal environments and oxidized biomass more than 3.8 billion years ago (Dodd 2017 ^PMID: 28252057^). Day length at the time of origin of very primitive photosynthesis is estimated to be around 12 h, while day length at the dawn of eukaryotes is estimated to be less than 18 h (Zahnle 1987; Sonett 1996 ^PMID: 8688061^; Williams 2000; Bartlett 2016) (Figure 3 and Supplementary File S4).

**Figure 3.**
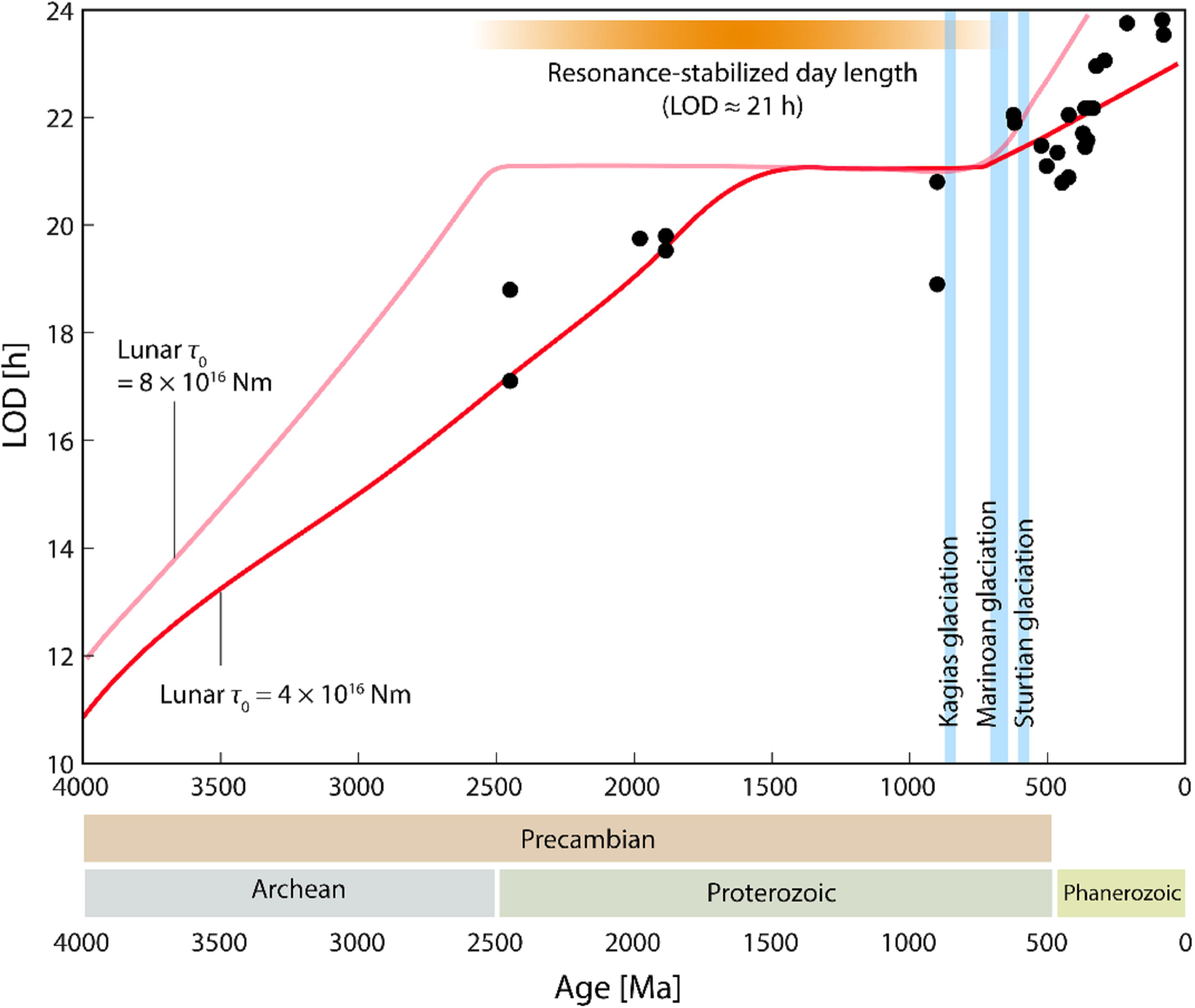
Changes of LOD over a period of 4000 million years (Ma). Black dots represent empirically determined LOD values based given by Williams. The two red functions represent different predictions of the long-term LOD changes by Bartlett and Stevenson based on varying choices of τ_0_, the lunar torque. Only the prediction based on the upper and lower range of τ_0_ values used for the simulation are shown for clarity.

Interestingly, our systematic investigation of mouse gene phylogenies revealed that genes exhibiting ultradian oscillatory patterns of expression are enriched of ancient genes, which arose at the early stages of life evolution. Many eukaryote species retain one (or more) copies of such genes; consistently, they have been classified as “essential” for mouse survival. Ultradian period oscillating genes comprised in mouse gene subsets are particularly enriched for the Unfolded Protein Response, *(Ddit3, Insig1, Dnajb9, Edem3, Hspa1b, Hspa5, Hspa13*) as well as the Protein Ubiquitination pathway *(Usp28, Usp46, Uspl1, Ube1dc1, Ube1l2, Ube2g2, Anapc10, Dnajb9, Hspa5, Psmc5, Anapc4, Dnajb1, Vhl)*.

Ultradian rhythms are known to play a key role in physiology, as evidenced by the 12-hour activation rhythm of the lRE1⍺-XBP1 pathway in the endoplasmic reticulum driving the two-peak activation of the Unfolded Protein Response (UPR)-regulated genes hallmarking hepatic lipid metabolism (Cretenet 2010 PMID:20074527) UPR is adaptive response conserved across kingdoms, phyla and species and committed to manage the accumulation of unfolded proteins in the endoplasmic reticulum to increase expression of genes entailed in endoplasmic reticulum function to re-establish/augment the capability of folding and modify proteins (Hollien 2013 ^PMID: 23369734^; lwata2012 ^PMID: 22796463^; Zhang 2016 ^PMID: 27256815^). On its side, the Ubiquitin/Proteasome interactive pathway is evolutionary conserved and essential in organisms ranging from yeast to mammals for the coordinated and temporal targeted degradation of short-lived proteins or proteins that do not fold accurately within the endoplasmic reticulum. Monoubiquitination is involved in endocytosis, subcellular protein localization/trafficking variations and DNA damage response, whilst multiple ubiquitination rounds play a role in protein targeting for proteasomal degradation (Amm 2014; Budhidarmo 2012)

The preferential usage among synonymous codons is a common property of many genomes from Prokaryotes to Eukaryotes. This bias, CUB, is thought to reflect a balance between mutational pressure and selective forces The differential SCU among genes would act as a regulator of the translational process, by shifting from a rapid peptide synthesis to a slower one in which the peptide chain has an appropriate time to fold correctly [Yu 2015 ^PMID:26321254^]. In simple model organisms, efforts were focused on the study of the relationship between SCU and translational efficiency/accuracy. For example, in the cyanobacterium *Synecocchus elongatus*, two strains with optimized and non-optimized (wild type) SCU for the KaiABC core clock system were tested for rhythmicity preservation. The wild type strain with a lower CUB was found to be more efficient in maintaining rhythmicity in different environmental conditions [Xu 2013 PMID: 23417065]. A non-optimal SCU was also tested in the Frequency (FRQ) circadian protein in *Neurospora crassa* through construction of an optimized version of the FRQ coding sequence: circadian rhythm obliteration was observed *in vivo* [Zhou 2013 PMID: 23417067; Zhou 2015 PMID: 26032251]. The usage of “unpreferred” synonymous codons was shown to slow-down FRQ mRNA translation, ensuring appropriate protein folding, especially for the Intrinsically Disordered Regions (‘IDP’) within it [Zhou 2013 PMID: 23417067; Zhou 2015 PMID: 26032251]. Studies on the *Drosophila melanogaster* circadian gene Period and CUB confirmed that artificially induced codon optimization would cause protein instability and alteration of circadian rhythmicity regulation [Fu 2016 PMID: 27542830]. Our analyses performed on mouse oscillating genes showed that ultradian genes generally have more relaxed codon usage. This phenomenon could be linked to the need for appropriate protein folding, although it is very hard to demonstrate this hypothesis for such a large gene set using computational modeling and functional assays.

The results of our study are globally in agreement with the general notion that ancient genes have a propensity for non-biased codon usage, as detected by a recent analysis on the human genome showing that gene age has a significant inverse correlation with SCU bias (with older genes having Codon Deviation Coefficient values toward 0) [Yin 2016, ^PMID:27609935^]. This issue is further complicated when gene expression is taken into account. Flowever, while housekeeping (ubiquitous, highly expressed) genes show optimized SCU [Ma 2014 PMID ^25011537^], selection for translational efficiency could be less relevant in case of tissue-specific genes or genes with highly variable expression, such as oscillating genes. However, we cannot exclude that, for some specific oscillatory gene, CUB could be determined by mechanisms resembling those found for circadian genes in *Neurospora Crassa* or *Synecocchus elongatus*.

Dawn/dusk transitions impact rhythmic gene expression as evidenced by experiments performed in *Neurospora Crassa* and showing that the greater part of oscillating transcripts managing metabolic functions are produced by dawn and dusk specific transcription (Sancar 2015 ^PMID:25762222^). In addition to diurnal patterns of solar illumination, Earth’s rotation on its axis dictates environmental temperature fluctuations. Life forms are accustomed to temperatures from the freezing to over the boiling point of water, but heat stress caused by temperatures barely exceeding the relative most favorable growth temperature severely limits survival (Richter 2010 ^PMID:20965420^). Interestingly, some of the oscillating genes comprised in the 12-hour periodicity subset *(Hspa1b, Hspa5, Hspa13, Dnajb1, Dnaja4, Dnajb9)* encode heat shock proteins (HSP). HSP work as molecular chaperones to avert the formation of anomalous protein aggregates and support proteins in the acquisition of their native structures, enriching an ancient signaling pathway activated by abrupt temperature increase (heat stress) harming vital cellular structures and obstructing crucial functions (Conway de Macario 2017 ^PMID: 28119916^; Carra 2017 ^PMID: 28364346^). Moreover, UPR activity fluctuates with ultradian pattern in mouse liver cells and especially in the endoplasmic reticulum. Interestingly, the expression of XBP1, driving one of the three UPR effector pathways, oscillates with 12-hour periodicity [Cretenet 2010 ^PMID:20074527^] and regulates the downstream expression of genes implicated in protein folding, turnover and proteo-toxicity control [Chaix 2014 ^PMID: 27738003^]. The presence of XBP1 binding sites on upstream sequences of 12h oscillating genes would support the hypothesis that these ultradian genes played a key role during the evolution of all life forms on planet Earth.

In addition to the genes oscillating with a 12-hour period, the 8h oscillating gene subset included ancient genes and enriched pathways related to sterol synthesis and oxidative stress. Fascinatingly, sterol biosynthesis is an O_2_-intensive process, and relevant molecular fossils contain steroidal hydrocarbons (steranes), which are the diagenetic remains of sterol lipids and hint at Paleoproterozoic sterol biosynthesis (Gold 2017 ^PMID:28264195\^. More elementary sterols can be synthesized by some bacteria, and complex sterols with modified side chains are synthesized by eukaryotes. Complex steranes found in ancient rocks suggest aerobic metabolic processes rather than the presence of eukaryotes, dating the divergence time of bacterial and eukaryal sterol biosynthesis genes around 2.31 billion years ago. This is consistent with the most recent geochemical evidence for the great oxidation event (Gold 2017 ^PMID: 28264195^). O_2_ production totally transformed Earth’s surface and atmosphere during the Proterozoic Eon, leading to substantial atmospheric O_2_ levels rise about 3 billion years ago with fading of the reducing conditions prevalent during the Archean. The great oxygenation event was hallmarked by considerable and irreversible O_2_ accumulation in the atmosphere starting in the early Paleoproterozoic and boosting during the Mesoproterozoic (Bekker 2004 ^PMID: 14712267^_j_ Cavalier-Smith 2006 ^PMID: 16754610^; Parnell 2010 ^PMID: 21068840^; Crowe 2013 ^PMID: 24067713^; Hamilton 2016 ^PMID:26549614^; Luo 2016 ^PMID:27386544^). Life exponentially increased during the great oxidation event, but oxygen is harmful for nucleic acids and avoiding exposure of these biomolecules to the reductive phases of the respiratory cycle could prevent oxidative damage. Possibly, the oldest unicellular autotrophic organisms, for instance the cyanobacteria, evolved ultradian rhythmicity to coordinate DNA replication and transcription with mitochondrial redox activity, differently from the more recent life forms, such as unicellular algae, hallmarked by circadian rhythmicity. Intriguingly, aerobic metabolism was more proficient, but O2 reactivity and toxicity threatened living organisms, which developed biochemical tools to defend against oxidative damage. These comprise enzymatic and non-enzymatic antioxidants blocking dangerous reactive oxygen species and among the genes oscillating with 8-hour periodicity we found several genes, such as *Pon1, Cat, Apoa1, Apoa2, Prdx1, Hmox2*, encoding proteins involved in redox reactions and other molecular interactions directly associated to antioxidant activity (Neumann 2003 ^PMID: 12891360^; Oda 2001 ^PMID: 11327831^; Larcher 2015 ^PMID: 26163000^; Barter 2004 ^PMID: 15486323^). Among these genes, particularly interesting is the *Cat* gene, encoding catalase, a highly conserved heme enzyme (Zámocký 2008 PMID. 18498226), capable hydrogene peroxide (H_2_O_2_) and present in the peroxisomes of nearly all aerobic cells; accordingly, the catalase evolutionary process started in the Archaean and boosted in the Proterozoic, in line with the increase in atmospheric O_2_ (Zámocký 2012 ^PMID: 22330759^; Zámocký 2014 ^PMID: 24846396^; Zámocký 2015 ^PMID: 25575902^).

## Conclusions

The expression of oscillating genes shows rhythmicity in more than one time domain and the harmonics of transcription periodicity could depend, further than on transcriptional dynamics, also on phylogenetic components related to geochemical changes and evolutionary mechanisms. Ultradian rhythms are crucial for organismal physiology and characterize essential biological functions and adaptative processes conserved across kingdoms, phyla and species, such as metabolic cycles and the response to cell stress. In particular, 12h oscillating genes are more likely to be ancient and essential, suggesting that these genes may have evolved to deal with the twice-daily cycles of transition between light/heat and dark/cold. Our comprehensive computational analyses show that the genes oscillating at the second and third harmonic of circadian rhythmicity are more phylogenetically conserved across all kingdoms of life and may represent an evolutionary genomic footprint.

## METHODS

### Primary dataset

We exploited a dataset compiled as previously described (Hughes 2009 ^PMID:19343201^) and available from GEO (GSE11923). Rhythmic genes were identified in mouse hepatic tissue using both COSOPT and Fisher’s G-test at a false-discovery rate of < 0.05. Clusters of rhythmic genes with period lengths of approximately 24h (> 20 and < 30 hours), 12h (> 10 and < 14 hours) and 8h (> 7 and < 9 hours) were observed. We selected lists of array probe IDs/nucleotide sequences for which oscillating expression levels were determined and performed our analyses on three oscillating gene subsets: 56 (8h), 202 (12h) and 2396 (24h) Ensembl Transcript IDs were retrieved from the primary dataset, by using BioDBnet (https://biodbnetabcc.ncifcrf.gov/db/db2db.php) db conversion tool, respectively.

### Analysis of gene genealogies

Gene phylogenies can be resolved by numerous computational techniques. Given the elevate number of investigated genes (more than 2000 oscillating genes), we opted to consider pre-computed phylogenetic gene trees from three popular bioinformatic resources, PhylomeDb (http://phvlomedb.org/). NCBI HomoloGene (http://www.ncbi.nlm.nih.gov/homologene/) and the OGEEv2 Portal (http://ogee.medgenius.info/).

PhylomeDB was programmatically accessed (Spring 2016) through the web resource, focusing on the Reference Model Species Metaphylome (id: 504) of *Mus musculus* and extracting from it data consisting of: the submitted mouse gene Ensembl ID (Ensembl Genes 83 release, January 2016; mouse genome version GRCm38.p4), official gene symbol and taxon name for the largest taxonomic group that comprises the investigated gene.

Secondarily, we extracted gene phylogenetic information from NCBI HomoloGene (build 146, accessed January 2016) using *“Mus musculus* [ORGN]” as query term. In particular, we manipulated the NCBI “summary output” consisting of HomoloGene Accession Number, gene-specific degree of conservation (resumed with the formula “Conserved in taxon XX”) and relative gene symbol.

From the OGEEv2 databank (accessed January 2017), we downloaded the “gene essentiality” dataset, a collection of tested essential and non-essential genes in multiple species, and “gene age”, an additional dataset for gene evolutionary properties (Supplementary Tables S1 and S2).

### Synonymous codon usage analysis in Mus musculus genome

Genomic investigations in a large spectrum of organisms have shown that the usage of synonymous codons is variable across species and genes. Several researchers demonstrated that asymmetry in the usage of codon within a certain codon family derives from the counterbalance of mutational pressure, selection and random drift. The quantification of the bias in synonymous codon usage, SCU (‘codon usage bias’ or ‘CUB’) can be performed by implementing numerous methods [Behura 2013 ^PMID:22889422^]

We primarily opted to calculate the “effective number of codons” (“ENC” or 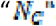) as a measure of the bias that, similarly to the “effective number of alleles“, that is based on 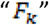 (average homozygosity of one degenerate codon family with “k” synonymous codons), while integer numbers are relative to the twenty codon family (2 is associated to methionine and tryptophan, that are encoded by one single codon) [Wright 1990 ^PMID: 2110097^].

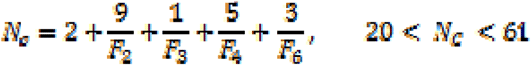

With some adjustments, ENC ranges from 20 (maximum bias with a single synonymous codon used for each amino acid) to 61 (minimum bias, with all possible codons used). Importantly, this index is not based on any knowledge about organism-specific highly expressed genes and this is quite useful especially for species in which gene expression levels are poorly known.

ENC was calculated by using the popular CodonW Package (http://codonw.sourceforge.net/, ref) for 26814 protein-coding FASTA sequences as retrieved by Ensembl Biomart (Ensembl v82, mouse genome: GRCm38.p4).

Secondarily, we considered another CUB index, the “Codon Adaptation Index” (CAI) (Sharp 1987, ^PMID: 3547335^) defined as “the geometric mean of the relative adaptiveness values (ω) of individual codons”:

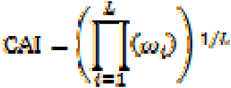

where “L” represents the total number of codons, while ω_i_ is the ratio of the observed codon frequency over the frequency of the most used codon for a certain amino acid. CAI ranges from 0 (usage of non-optimized codons) to 1 (usage of optimized codons). The “most used” or “most abundant” codons are calculated on codon usage of highly expressed genes. Thus, this measure necessitates of the knowledge of constitutive/highly expressed genes for a certain organism. Focusing on the mouse genome, we relied on the JCat webtool (http://icat.de, ref) that calculates CAI for a series of prokaryotic genomes and a limited number of Eukaryotes, among which *Mus musculus* (run with default parameters, accessed at January 2016). We used the same 26814 FASTA sequences as input.

Distributions of ENC/CAI values for the entire mouse genome and the oscillatory gene subsets were compared by Wilcoxon rank sum tests. Furthermore, in order to “minimize” the impact of mutational pressure in determining CUB, we considered only the sequences with a comparable base composition. In detail, we calculated the GC content at third codon positions (‘GC_3s_’) for each mouse sequences and discarded those with GC content within the first or the third quartile. We then compared ENC or CAI distributions for the retained sequences.

### Relative Enrichment analysis for known TFBS in oscillating genes

We performed a differential enrichment analysis for known transcription binding, by focusing on the 5000 bp at the 5’ end of CDS initiation site for the classes of oscillating genes and, as controls, five sample datasets of 1500 randomly chosen upstream regions from mouse genes and additional five samples of 500 sequences. Sequences were retrieved from Ensembl build 84, by querying the GRCm38p2 mouse genome version through the Biomart tool. Background sequences were randomly picked up from the entire dataset (23111 regions), with the exclusion of those related to oscillating genes.

AME good usage practices (http://meme-suite.org/doc/ame.html?mantype=web) suggest that sequences should have comparable lengths, so we opted to work with a constant region size (5000 bp) for all the considered genes. We carried out 18 distinct runs of the AME program (MEME suite, 4.11.4 version) (Bailey 2009 ^PMID: 19458158^) as follows: upstream sequences of 8h oscillating genes versus the five control datasets; the same approach for 12h and 24h oscillating genes sequences; 8h oscillating genes versus 12h oscillating genes (using the second dataset as control); 12h oscillating genes versus 24h oscillating genes; 8h oscillating genes versus 24h oscillating genes.

The procedure was repeated also for the 500 sequences control datasets. Three datasets of “meme-formatted” mouse regulatory motifs were screened, for a total of 825 elements.

AME output is produced in the form of textual and html files in which relevant enriched motifs are sorted according Fisher’s Exact test p-values and adjusted p-values. Command lines and raw output are present in Supplementary File S3.

### Eukaryotic protein age analyses

Protein age was evaluated by ProteinHistorian, a software based on three inputs, a species tree, a protein family database, and an ancestral family reconstruction algorithm with two parsimony algorithms: Dollo parsimony and Wagner parsimony. Dollo parsimony is based on the assumption that gaining a complex structure is much rarer than losing one. Thus, it assumes that there was a single gain event for each family, potentially followed by many losses in specific lineages. In other words, under Dollo parsimony, a family’s origin is the most recent common ancestor (MRCA) of all species in which it is observed. Wagner parsimony allows multiple gain and loss events in an ancestral family reconstruction, as well as the ability to set weights on the relative likelihood of these events. By default, ProteinHistorian uses a relative gain penalty of 1. Since we focused on eukaryotic species in which horizontal gene transfer is rare, this largely serves to prevent false positives in the protein family databases from biasing age distributions. (Capra 2012 ^PMID: 22761559^).

(http://lighthouse.ucsf.edu/ProteinHistorian/)

## ACKNOWLEDGMENTS

Financial support: the study was supported by the “5×1000” voluntary contribution and by a grant to GM through Division of Internal Medicine and Chronobiology Unit (RC1403ME50, RC1504ME53, RC1603ME43 and RC1703ME43), IRCCS Scientific Institute and Regional General Hospital “Casa Sollievo della Sofferenza”, Opera di Padre Pio da Pietrelcina, San Giovanni Rotondo (FG), Italy. AR and NG, were funded by the German Federal Ministry of Education and Research (BMBF)—eBio-CIRSPLICE - FKZ031A316. NG was additionally funded by the Berlin School of Integrative Oncology (BSIO) of the Charité – Universitätsmedizin Berlin.

## AUTHOR CONTRIBUTIONS

G.M. conceived the study; T.M., A.R., N.G., J.H., C.F., D.C., T.B. and S.C. performed the bioinformatics analysis; F.S. computed length of day across centuries; G.M., N.G. and A.R. provided funding for the study; G.M., S.C., A.R., N.G., J.H. wrote the paper; G.M.,T.M., A.R., N.G., C.F., F.S., D.C., T.B., S.C. and J.H. approved the final version of the manuscript.

## MATERIALS & CORRESPONDENCE

Gianluigi Mazzoccoli (Division of Internal Medicine and Chronobiology Unit, IRCCS “Casa Sollievo della Sofferenza”, 71013 San Giovanni Rotondo (FG), Italy.

## Supplementary File S4

### Long-term changes of Earth’s rotation period (length-of-day)

The assessment of long-term changes of day length on Earth is a matter of investigation since many decades. While it was first noticed that the length-of-day (LOD) increased during Earth’s history, in 1987 Zahnle and Walker came to the conclusion that there are good arguments that in the Precambrian era, when the deceleration of the LOD would have reached about 21 h, the period would have been resonant with the semidiurnal atmospheric tidal torque (10.5 h). At this time, a stabilizing effect on the LOD would have occurred since the atmospheric tidal torque would have had the same magnitude (but opposite sign) as the lunar oceanic torque. This resonance condition would have stabilized the LOD at a value of about 21 h. Recently, Bartlett and Stevenson came to the conclusion that this resonance condition would have broken by sudden atmospheric temperature increases like the deglaciation happened at the end of the Precambrian (i.e. the Marinoan and Sturtian glaciations), causing anfurther deceleration of the LOD to the current value of 24 hours. Figure 1 shows the prediction of LOD variations based on the simulation performed by Bartlett and Stevenson. Empirically determined values of LOD were added according to the publication of Williams.

